# Serotonergic signaling plays a deeply conserved role in improving oocyte quality

**DOI:** 10.1101/2023.03.22.533887

**Authors:** Erin Z. Aprison, Svetlana Dzitoyeva, Ilya Ruvinsky

## Abstract

Declining germline quality is a major cause of reproductive senescence. Potential remedies could be found by studying regulatory pathways that promote germline quality. Several lines of evidence, including a *C. elegans* male pheromone ascr#10 that counteracts the effects of germline aging in hermaphrodites, suggest that the nervous system plays an important role in regulating germline quality. Inspired by the fact that serotonin mediates ascr#10 signaling, here we show that serotonin reuptake inhibitors recapitulate the effects of ascr#10 on the germline and promote healthy oocyte aging in *C. elegans*. Surprisingly, we found that pharmacological increase of serotonin signaling stimulates several developmental processes in *D. melanogaster*, including improved oocyte quality, although underlying mechanisms appear to be different between worms and flies. Our results reveal a plausibly conserved role for serotonin in maintaining germline quality and identify a class of therapeutic interventions using available compounds that could efficiently forestall reproductive aging.

## Introduction

In animals, reproductive costs can be staggering. For example, *C. elegans* hermaphrodites produce ∼100 embryos per day (McMullen et al., 2012), a biomass equivalent to the body mass of the mother! Because animals balance the demands of reproduction against activity, growth, and maintenance (Pontzer and McGrosky, 2022), elaborate regulatory mechanisms must ensure that gamete production, including stem cell proliferation and their subsequent differentiation, is well matched to the external conditions throughout the reproductive span. Studies over the past several decades revealed proximal mechanisms that regulate stem cell division, particularly those related to signaling from the niche (Morrison and Spradling, 2008). However, to match germline production to specific pressures imposed by the environment almost certainly requires coordinated signaling by the nervous system. A better understanding of these largely unknown mechanisms may open new avenues for manipulating the germline.

Sex pheromones are a useful tool for investigating the ways in which the nervous system regulates the germline. These small-molecule signals exchanged by potential mating partners are detected by sensory neurons and can ultimately alter reproductive physiology. For example, the major male pheromone in *C. elegans*, ascr#10 (Izrayelit et al., 2012), increases the number of germline precursor cells in adult hermaphrodites (Aprison and Ruvinsky, 2016), acting via a particular serotonergic neuronal circuit that relies on the MOD-1 receptor (Aprison et al., 2022b; Aprison and Ruvinsky, 2019a, b, 2022). The principal effect of ascr#10 on the hermaphrodite germline is to increase proliferation of mitotic progenitors (Aprison et al., 2022a) in adults after the onset of reproduction (Aprison and Ruvinsky, 2019b), that is, well after the irreversible switch from sperm to oocyte production (Kimble and Crittenden, 2007). One downstream consequence of this increase is a greater number of germline precursor cells, while another is the improved quality of the oogenic germline, particularly in older hermaphrodites (Aprison et al., 2022a). Oocytes of the ascr#10-exposed hermaphrodites benefit from a more youthful morphological appearance, decreased rate of chromosomal nondisjunction, and reduced embryonic lethality in the progeny both in the wild type and in genetically sensitized backgrounds.

Reduced oocyte quality is the major driver of reproductive decline. For example, maternal age is a major risk factor for aneuploidy (Nagaoka et al., 2012) and women at 45 years of age have few oocytes that could support embryonic development (Centers for Disease Control and Prevention, 2018). Similar defects – production of morphologically defective oocytes, increased rates of chromosomal nondisjunction, as well as depleted stores of mitotic germline precursors – are seen in the germlines of the two preeminent invertebrate model organisms, *C. elegans* and *D. melanogaster* (Andux and Ellis, 2008; Garigan et al., 2002; Hughes et al., 2007; Ishibashi et al., 2020; Kocsisova et al., 2019; Luo et al., 2010; Miller et al., 2014; Pickett et al., 2013; Qin and Hubbard, 2015; Zhao et al., 2008).

Identifying specific signaling pathways involved in germline quality maintenance may lead to the development of remedies that support healthy reproductive aging (Achache et al., 2021; Templeman et al., 2018). Our previous studies revealed that ascr#10 can improve oocyte quality (Aprison et al., 2022a) and that serotonergic signaling is necessary (Aprison and Ruvinsky, 2019a) and sufficient (Aprison et al., 2022b) to recapitulate several ascr#10 effects. We therefore wondered whether compounds that increase serotonergic signaling could improve quality of the oogenic germline in *C. elegans*. We also tested whether stimulation of serotonin signaling could promote germline health in a distantly related species.

## Results & Discussion

### Increased serotonin signaling promotes germline proliferation in *C. elegans* hermaphrodites

Exposure to ascr#10 increases serotonin signaling from NSM and HSN neurons in reproductive *C. elegans* hermaphrodites, whereas loss of serotonin signaling blocks ascr#10 effects on the germline (Aprison and Ruvinsky, 2019a, b). Moreover, mutations in the serotonin transporter gene *mod-5* that effectively increase serotonin signaling because they reduce reuptake of the neurotransmitter at the synapse (Ranganathan et al., 2001), show greater germline proliferation (Aprison et al., 2022b). For these reasons, we tested whether pharmacological increase of serotonergic signaling can recapitulate the ascr#10 effects on the germline.

Selective serotonin reuptake inhibitors (SSRIs) constitute a large class of commonly prescribed medications that promote serotonergic signaling. An SSRI fluoxetine (Prozac, etc.) increased germline proliferation (Figure 1A) demonstrating that pharmacological stimulation of serotonergic signaling is sufficient to recapitulate at least this aspect of ascr#10 effect on the hermaphrodite germline. Previously we demonstrated that increased supply of germline precursors due to increased proliferation on ascr#10 leads to increased physiological cell death during prophase of Meiosis I (Aprison et al., 2022a). Similarly, we found that fluoxetine treatment resulted in increased cell death (Figure 1B).

**Figure 1.**
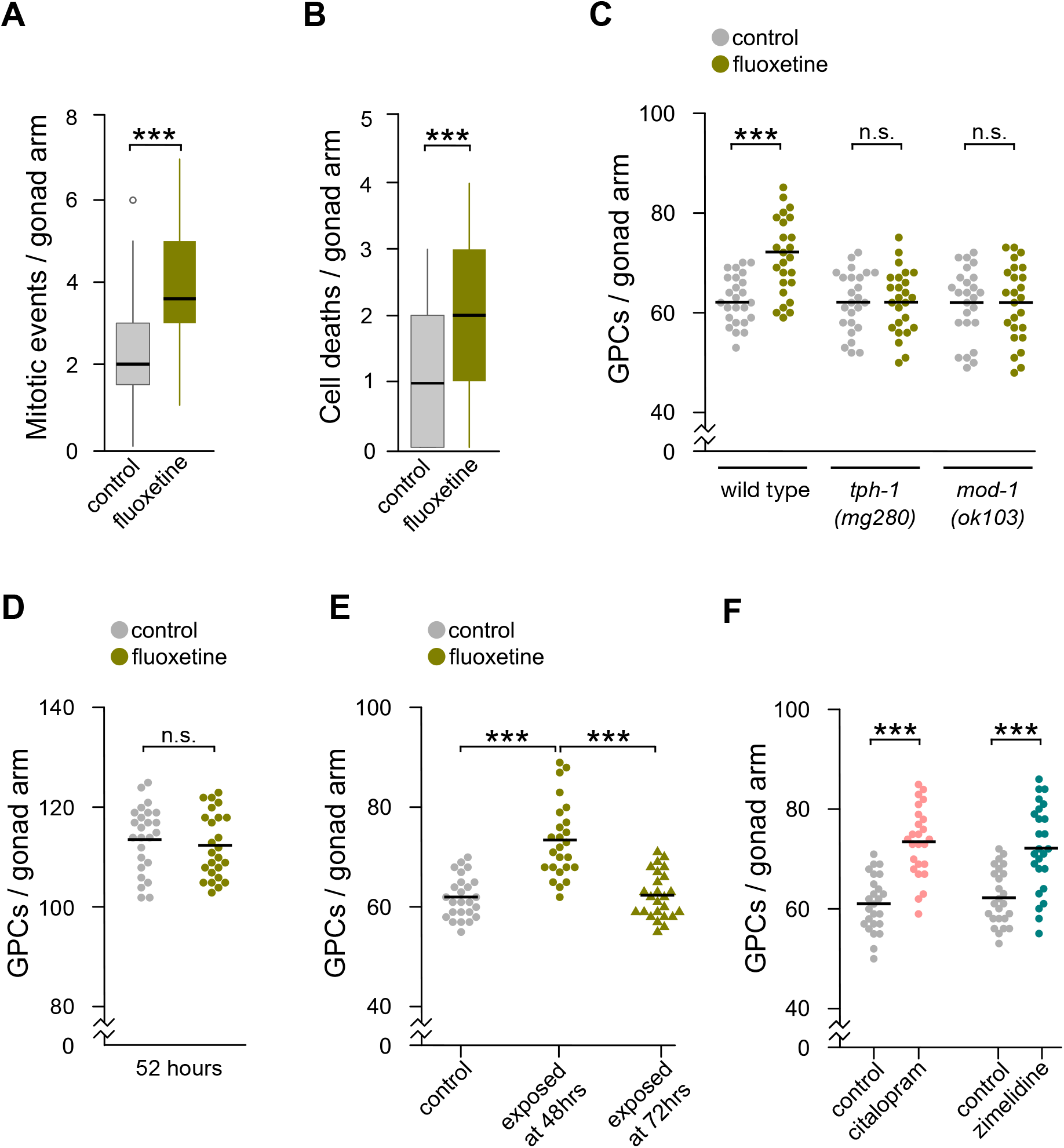
Effects of increased serotonergic signaling on the oogenic germline in *C. elegans*. **(A)** Mitotic events and **(B)** dying cells in the gonads of Day 3 adult hermaphrodites on fluoxetine vs. untreated controls. **(C)** GPCs in N2 wild type, *tph-1(mg280)*, and *mod-1(ok103)* Day 5 adult hermaphrodites on fluoxetine vs. untreated controls. GPCs in N2 wild type hermaphrodites exposed to fluoxetine from **(D)** hatching until 52 h (adult, just before onset of egg laying) or **(E)** indicated ages until Day 5 of adulthood vs. untreated controls. **(F)** GPCs in Day 5 adult N2 wild type hermaphrodites on SSRIs citalopram and zimelidine vs. untreated controls. In **C, D, E**, and **F**, each dot represents the number of GPCs in an individual hermaphrodite. Black bars denote means. ***, p<0.001. See Table S1 for primary data and details of statistical analyses.

As hermaphrodites age, they experience progressive loss of germline precursors (Andux and Ellis, 2008; Garigan et al., 2002; Hughes et al., 2007; Ishibashi et al., 2020; Kocsisova et al., 2019; Luo et al., 2010; Miller et al., 2014; Pickett et al., 2013; Qin and Hubbard, 2015; Zhao et al., 2008). Here, as germline precursor cells (GPC) we considered nuclei in the Progenitor Zone (PZ), as previously defined (Crittenden et al., 2006). The PZ contains stem cells, cells that are finishing the last mitotic cycle, and those that have entered meiotic S-phase (Hubbard and Schedl, 2019). Fluoxetine increased the number of GPCs in aging hermaphrodites, in a manner dependent on tryptophan hydroxylase TPH-1, the rate-limiting enzyme of serotonin biosynthesis (Sze et al., 2000), and MOD-1, the only known serotonin receptor involved in regulation of the oogenic germline in response to ascr#10 (Aprison and Ruvinsky, 2019a, 2022) (Figure 1C).

Exposure to ascr#10 does not promote germline proliferation in hermaphrodites prior to the onset of egg laying (Aprison et al., 2022a; Aprison and Ruvinsky, 2019b) and exposure to this pheromone has to commence before the onset of egg laying to increase germline proliferation (Aprison et al., 2022a). Consistent with these temporal limitations on ascr#10 effects, we found that fluoxetine did not alter germline proliferation in adults prior to egg laying (Figure 1D) or if exposure started after the onset of reproduction (Figure 1E). Fluoxetine was not the only serotonin signaling-promoting drug that affected the germline. Hermaphrodites exposed to structurally distinct SSRIs – citalopram (Celexa, etc.) and zimelidine – also had more GPCs (Figure 1F).

### Fluoxetine improves oocyte quality in *C. elegans*

Because fluoxetine boosted progenitor cell proliferation in the PZ and the incidence of germline cell death, two processes required for germline quality improvement by ascr#10 (Aprison et al., 2022a), we asked whether fluoxetine could improve oocyte quality. The occurrence of developmental arrest in the offspring of a mating between young males and older hermaphrodites that have used up all of their self-sperm, is a reliable indicator of oocyte quality (Andux and Ellis, 2008; Hughes et al., 2007; Luo et al., 2010). Using this experimental set up, we observed lower incidence of developmental arrest in the offspring of Day 5 hermaphrodites that were aged in the presence of fluoxetine (Figure 2A). Similarly, fluoxetine lowered the fraction of arrested embryos produced via selfing by hermaphrodites on Days 4 & 5 of adulthood (from ∼5.3% to 3.8%; Table S1). The rate of nondisjunction (Figure 2B) and morphologically defective oocytes (Figure 2C, D) were also significantly reduced by fluoxetine, demonstrating that these effects of ascr#10 on the germline were recapitulated by fluoxetine just as increased proliferation and cell death (Figure 1).

**Figure 2.**
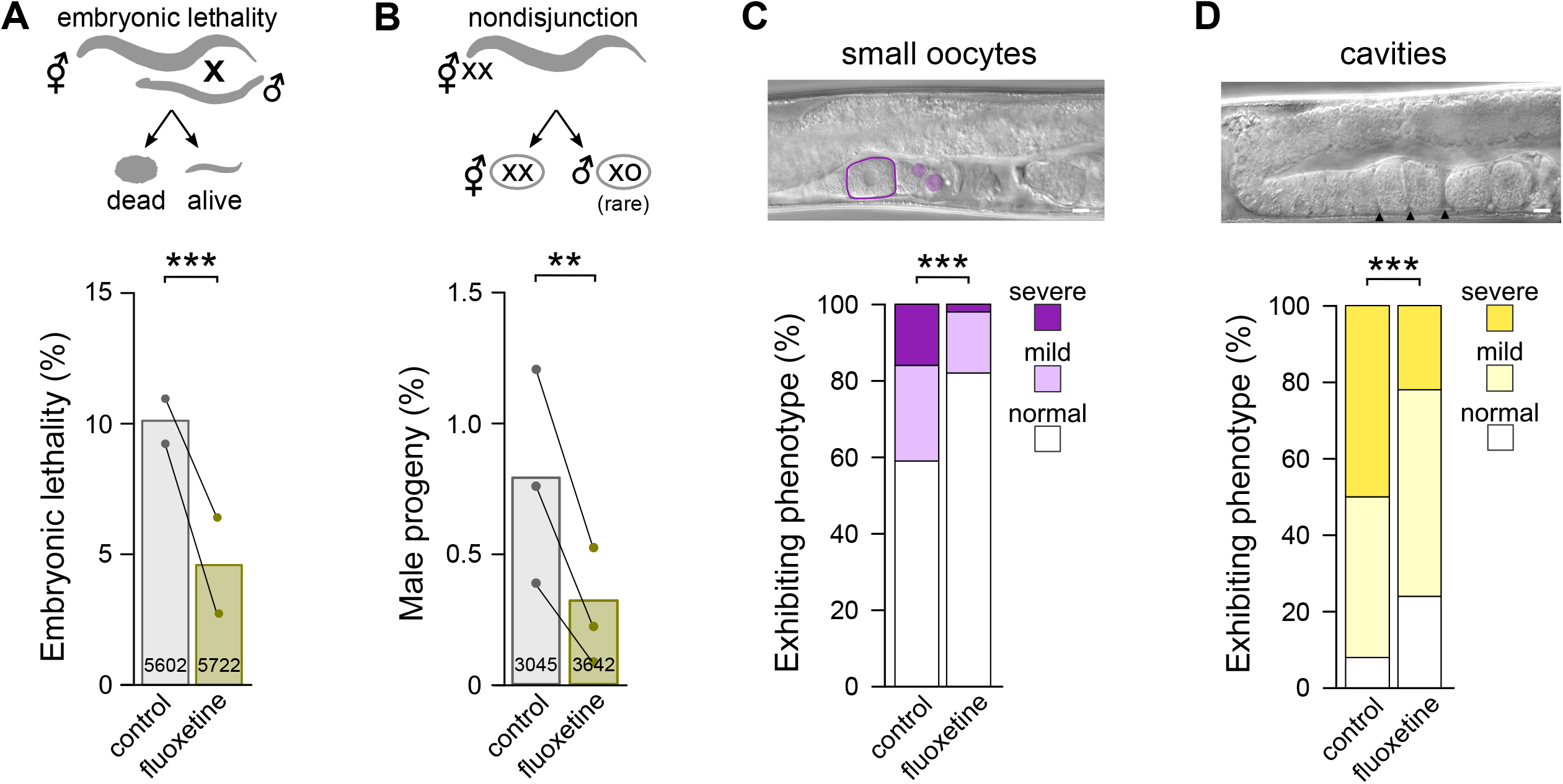
Fluoxetine improves oocyte quality in *C. elegans*. **(A)** Embryonic lethality in the progeny of Day 5 adult N2 wild type hermaphrodites aged on fluoxetine vs. untreated controls; both mated to young N2 males. **(B)** Incidence of males in the self-progeny of Day 5 adult N2 wild type hermaphrodites aged on fluoxetine vs. untreated controls. In **A** and **B**, each dot represents an individual experiment paired with its control. Progeny totals are shown inside bars. **(C)** Abnormally sized oocytes and **(D)** gaps between oocytes in Day 7 adult N2 hermaphrodites aged on fluoxetine vs. untreated controls. In **C** and **D** scale bars = 10 μm. **, p<0.01; ***, p<0.001. See Table S1 for primary data and details of statistical analyses.

### Fluoxetine alters multiple developmental processes in *D. melanogaster*

To test whether the beneficial effects of increased serotonergic signaling on the oogenic germline were restricted to *C. elegans*, we examined the germlines of fluoxetine-treated *D. melanogaster* females. We noted significant increase in the number of vitellogenic-stage egg chambers in females exposed to fluoxetine (Figure 3A). Unlike in *C. elegans*, where SSRIs promoted germline proliferation (Figure 1), egg chambers in fluoxetine-treated females showed no evidence of increased numbers of germline cells (provisional oocyte and nurse cells). Instead, we observed significantly increased proliferation of follicle cells, particularly in Stage 3 and 4 egg chambers (Figure 3B). During ovarian development, the population of follicle cells considerably expands between Stages 2 and 5, just prior to ceasing divisions around Stage 6 (King, 1970). Follicle cells form an epithelial layer that surrounds the germline cyst and generates the eggshell (King, 1970). Consistent with higher rates of proliferation, we found that egg chambers of flies exposed to fluoxetine contained more follicle cells (Figure 3C) and were significantly larger than in control flies (Figure 3D). However, sizes of terminal oocytes were equivalent between fluoxetine-treated and control flies (Figure 3E), implying the existence of a regulatory mechanism that ensures relative volume uniformity of mature oocytes.

**Figure 3.**
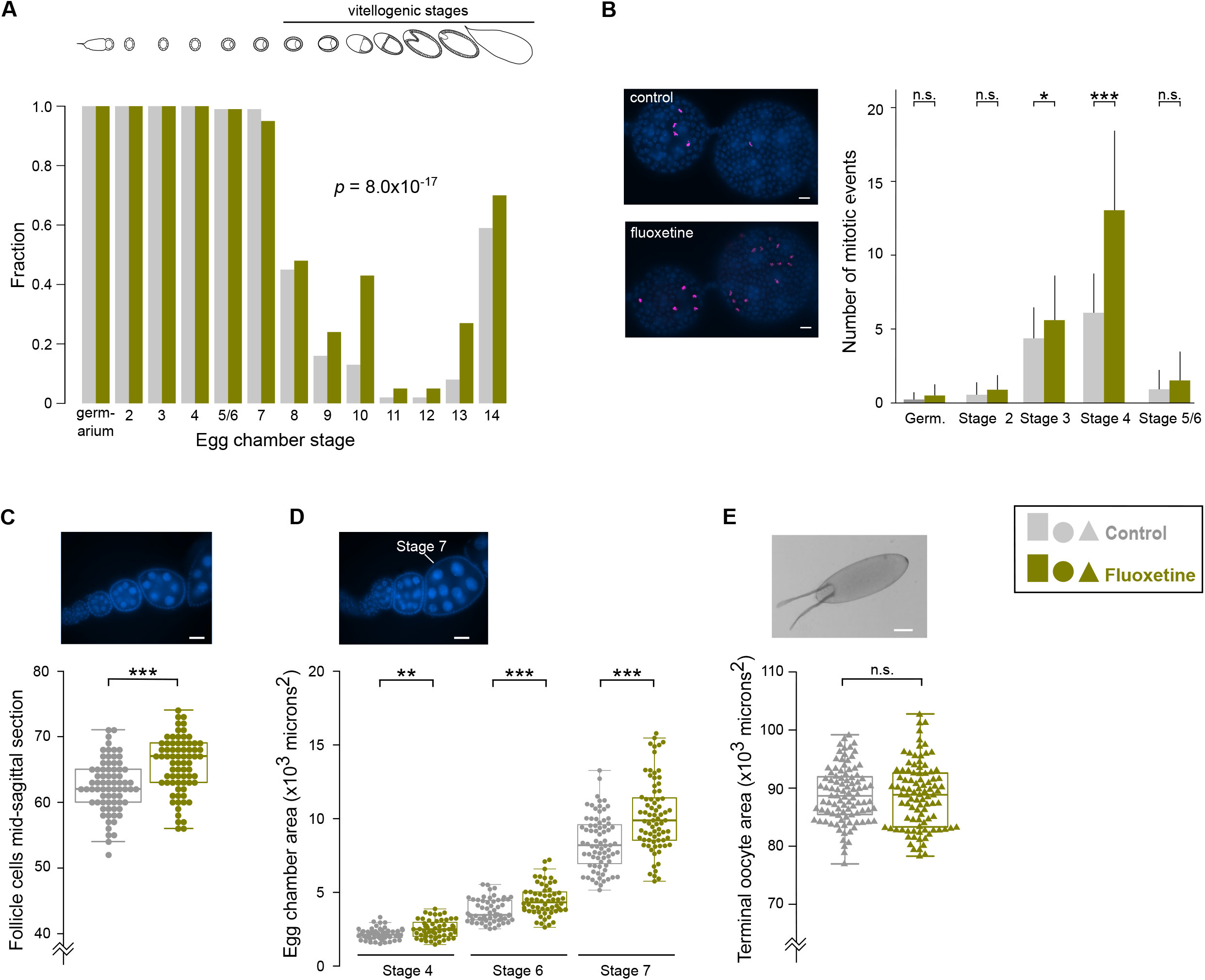
Effects of fluoxetine on ovaries in *D. melanogaster*. **(A)** The fraction of ovarioles that contain the designated egg chambers. These ovarioles were from mated Drosophila females raised on fluoxetine vs. untreated control. Ovarioles were dissected on Day 5 of adulthood. **(B)** Representative DAPI-stained images and quantification of mitotic events in egg chambers of Day 5 females raised on fluoxetine vs. untreated control. pH3-positive cells are shown in magenta. Scale bars = 10μm. In **B**, Germ. = germarium. **(C)** A representative DAPI-stained image and the number of follicle cells counted in mid-sagittal sections of Stage 7 egg chambers dissected from Day 5 mated females raised on fluoxetine vs. untreated control. Scale bar = 20μm. **(D)** Representative images of DAPI-stained egg chambers and quantification of their area from Day 5 mated females raised on fluoxetine vs untreated control. Scale bar = 20μm. **(E)** Area of terminal oocytes dissected from Day 5 mated females raised on fluoxetine vs. untreated control. Scale bar = 100μm. In **B** error bars denote standard deviation. In **C, D**, and **E**, horizontal lines represent means. *, p<0.05; **, p<0.01; ***, p<0.001. See Table S1 for primary data and details of statistical analyses.

Two points warrant additional comments. First, a previous study reported deleterious effects of fluoxetine exposure on the oogenic germline in Drosophila (Willard et al., 2006). We believe that the apparent discrepancy between that study and the results reported here is due to different fluoxetine concentrations. We applied approximately 17,000-fold less of the drug than the 5 mg/ml concentration used by Willard *et al*. Consistent with this notion, we saw no increase of follicle cell proliferation even when we increased fluoxetine concentration 10X from the usual 0.9*μ*M used throughout this study (Figure S1A). This result implies that the Drosophila reproductive system responds to low-intermediate fluoxetine concentrations, whereas much higher concentrations of this compound may be toxic.

Second, some but not all fluoxetine effects we observed appear to require mating. For example, in virgin flies fluoxetine did not increase the number of vitellogenic-stage egg chambers (Figure S1B) as it did in mated flies (Figure 3A). Mating alone also did not increase the number of late egg chambers (Figure S1C), indicating a synergistic effect of mating and increased serotonin signaling. Whereas in both mated (Figure 3D) and virgin flies (Figure S1D) exposure to fluoxetine enlarged early vitellogenic-stage egg chambers, only mated flies had more follicle cells on fluoxetine (compare Figure 3C and Figure S1E). Mating is known to alter multiple traits in Drosophila females (Avila et al., 2011; Hoshino and Niwa, 2021), and our results suggest that increased serotonergic signaling further modulates several reproductive traits beyond those affected by mating.

### Fluoxetine improves oocyte quality in older *D. melanogaster*

As a first step in testing whether fluoxetine altered reproductive aging in *D. melanogaster* females, we examined whether chronic exposure to fluoxetine over several weeks continued to affect the germline in ways comparable to the shorter exposure described above. We found that 30-day-old females continuously exposed to fluoxetine since achieving adulthood still had more mitotic events and greater numbers of follicle cells (Figure 4A, B) as well as larger egg chambers (Figure 4C) compared to untreated controls (compare to Figure 3B, C, D).

**Figure 4.**
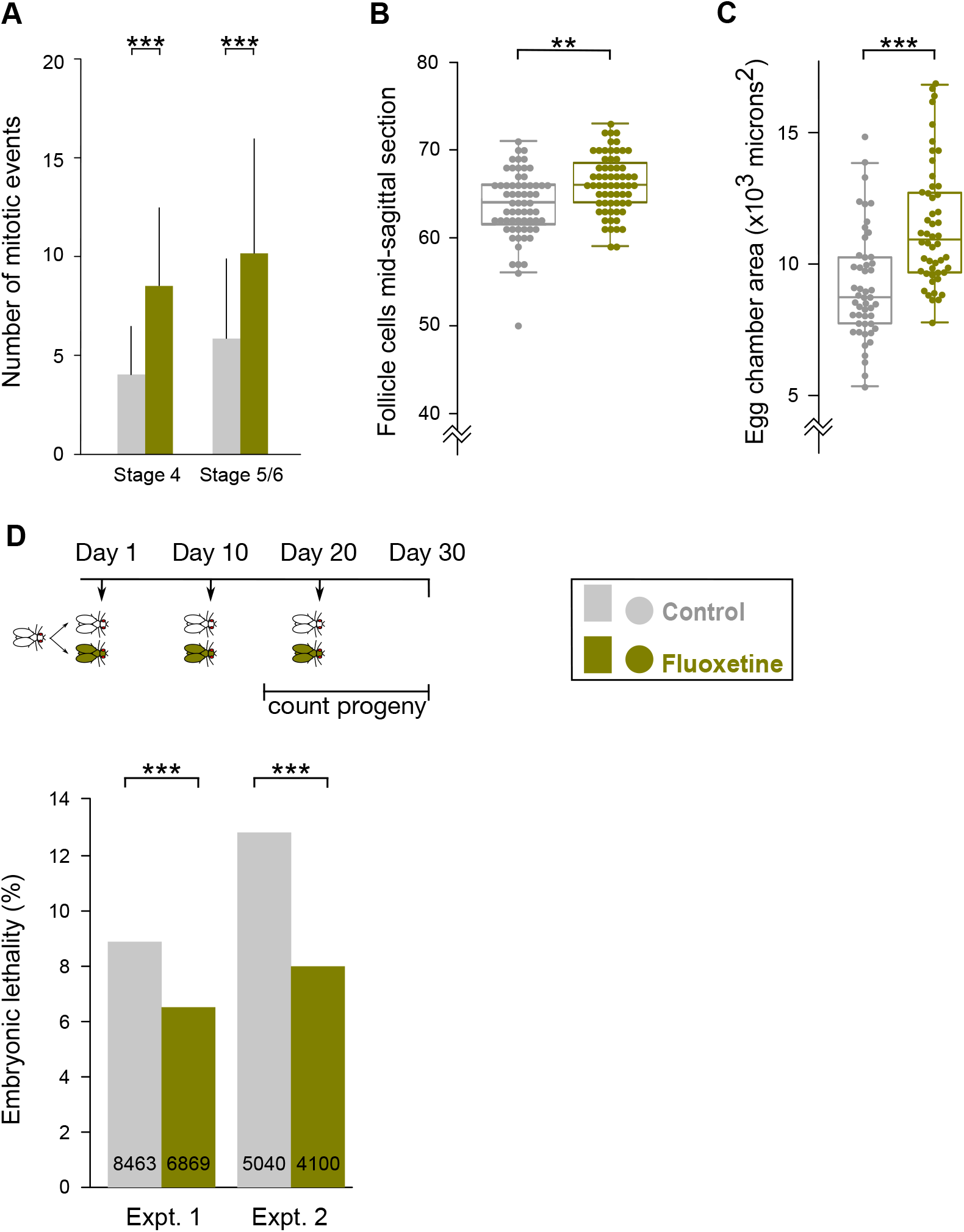
In aging Drosophila females, fluoxetine affects events in the ovaries and reduces embryonic lethality in offspring. **(A)** Mitotic events in egg chambers of Day 30 females raised on fluoxetine vs. untreated control. Bars indicate standard deviation. **(B)** Number of follicle cells surrounding mid-sagittal sections of Stage 7 egg chambers from Day 30 females raised on fluoxetine vs. untreated control. **(C)** The area of Stage 7 egg chambers from Day 30 females raised on fluoxetine vs. untreated control. **(D)** Schematic of experimental plan (down arrows indicating mating) and quantification of embryonic lethality in the offspring of mothers raised on fluoxetine vs. untreated control. Two independent experiments are shown. Embryonic lethality data pertain to offspring from matings 2 and 3 (Days 15 to 30, respectively). Numbers in bars are total eggs counted. **, p<0.01; ***, p<0.001. See Table S1 for primary data and details of statistical analyses.

To test whether exposure to fluoxetine ameliorated the effects of aging on oocyte quality in *D. melanogaster*, we mated individually-housed females to new young males every 10 days and monitored the resulting progeny. Offspring of older mothers that were maintained on fluoxetine suffered ∼20% lower embryonic lethality than the offspring of mothers under control conditions (Figure 4D). Therefore, in Drosophila, like in *C. elegans*, exposure to fluoxetine improved quality of the aging oogenic germline.

### Deep conservation of serotonin effects on oocyte quality

We emphasize two major findings of this study. First is that in both a nematode and an arthropod, increased serotonergic signaling alters multiple developmental events during oogenesis, although the specific targeted processes appear to be different. In *C. elegans*, the principal physiological effect of increased serotonergic signaling is greater proliferation of germline precursor cells, whereas in Drosophila, fluoxetine exposure increases proliferation of somatic follicle cells that surround the egg chamber. Despite these differences, it is possible however, that increased serotonin signaling in these two divergent animal lineages has the same ultimate effect. Increased GPC production in *C. elegans* leads to increased physiological cell death (Aprison et al., 2022a), a process that likely salvages organelles, nutrients, etc. to improve quality of the surviving oocytes (Tilly, 2001). A major function of follicle cells in Drosophila is to synthesize yolk proteins and thus to provision the oocyte (Brennan et al., 1982). Thus, in both worms and in flies elevated serotonin signaling induces changes that increase oocyte provision. Viewed in this way, reproductive effects of serotonin in worms and flies may reflect “deep homology” (Shubin et al., 2009). That is, in these distantly related lineages, elevated serotonin signaling induces different changes that nevertheless increase reproductive investment. Our work in *C. elegans* delineated the serotonergic neuronal circuit that regulates germline development (Aprison et al., 2022b; Aprison and Ruvinsky, 2019a, b, 2022). Discovering the structure, receptors, and signaling logic of the corresponding circuit(s) in Drosophila would be useful to further evaluate the hypothesis of “deep homology”.

Our second major finding is that pharmacological increase of serotonin signaling using commonly-prescribed generic medications improves the quality of the female germline in older *C. elegans* and *D. melanogaster*. We interpret the beneficial effects of SSRIs as a consequence of better oocyte provision, even if it is accomplished by different means in these two species. A major question for future work is whether drugs that increase serotonin signaling have similar effects on oocytes in other species.

Pheromones alter development without imposing high pleiotropic costs, typical of severe loss-of-function mutants often used in developmental biology, since they evolved to promote fitness. Our results advocate the study of pheromones as a powerful entry point for identifying pathways that regulate development in quantitative yet physiologically relevant ways. Moreover, the study of neuronal circuits that non-autonomously regulate development of other organs, including the reproductive system, promises to reveal unanticipated classes of therapeutic targets against disorders such as aging.

## Material and methods

### General

The comparison of volumes of embryos to mother is based on the WormBase-provided estimates of early embryo volume ∼2×10^4^ *μ*m^3^ and young adult volume ∼3×10^6^ *μ*m^3^.

The following drug concentrations were used in all *C. elegans* experiments – 1μM for fluoxetine, 10μM for citalopram, and 50μM for zimelidine. All *D. melanogaster* experiments, except for one shown in Figure S1A, were done using 0.9μM fluoxetine. The fluoxetine concentrations used in this study are comparable to those seen in the plasma of human patients (Amsterdam et al., 1997).

All slides were imaged on a Leica DM5000B microscope using a Retiga 2000R camera. ImageJ was used to measure objects in micrographs.

### Strains and maintenance

*Caenorhabditis elegans –* N2 wild type, MT15434 *tph-1(mg280)*, MT9668 *mod-1(ok103)*. All worm strains were obtained from the Caenorhabditis Genetics Center and were maintained at 20^°^C on OP50 under standard nematode growth conditions (Brenner, 1974). Synchronized populations were obtained using alkaline hypochlorite treatment of gravid hermaphrodites. Eggs isolated by this treatment were allowed to hatch overnight (no more than 16 hours) in M9 buffer with rotation at 20^°^C (Sulston and Hodgkin, 1988). The arrested L1 larvae were transferred onto lawn plates of OP50 the following morning at a density of 30-60 larvae per plate. Based on our experience staging N2 hermaphrodites, 48 hours after release from larval arrest was designated as Day 1 of adulthood (Aprison and Ruvinsky, 2014).

At this stage, worms are adults but have not yet begun to lay eggs. Some strains were slightly delayed in their development compared to the N2 wild type, so timing of experiments using these strains was adjusted to accommodate their developmental delay. On Day 1 of adulthood, hermaphrodites were transferred in populations of 30 worms per plate to either control or treatment plates. Adult hermaphrodites were moved to fresh plates every other day.

The wild type *Drosophila melanogaster* strain Oregon-R, a gift from Robert Holmgren, was used in all experiments. Flies were reared on Formula 4-24 Instant Drosophila Medium (Carolina Biological Supply Company, Burlington, NC 27216-6010) and maintained at 25^°^C.

### Conditioning *C. elegans* plates with pharmaceuticals

2 mg/ml stock solutions of fluoxetine hydrochloride (Sigma PHR1394), citalopram hydrobromide (Sigma PHR1640), and zimelidine dihydrochloride (Sigma Z101) were prepared in water and stored at 4^°^C. Fresh stocks were made every month. An appropriate amount of each stock was diluted and applied to plates to achieve the desired concentration based on a total volume of 10mL for 60mm plates and 3mL for 35mm plates. Plates were incubated at 20^°^C overnight to allow the drug to be absorbed.

Control plates were prepared in the same manner using water as the control. The following day the plates were seeded with 20μL of a 1:10 dilution of OP50 overnight culture and allowed to grow overnight at 20^°^C before use.

### *C. elegans* dissection and immunohistochemistry

Worm dissection and antibody staining were modified from (Crittenden et al., 2017). Hermaphrodites (60 – 90 per experiment) aged to the appropriate stage were picked into 200μL PBS-0.1%Tween 20 with 0.25mM levamisole in a large glass petri dish, and cut to extrude the germline. The dissected animals were transferred to a 1.5mL microcentrifuge tube and incubated with 3% paraformaldehyde in PBS-Tween for 30 minutes at 20^°^C with rocking. The paraformaldehyde was washed off and the worms were fixed in -20^°^C methanol for 30 minutes. The methanol was washed off and the worms were blocked with 3% bovine serum albumen in PBS-Tween for 30 minutes at 20^°^C with rocking. The blocking agent was washed off and the worms were incubated overnight with the primary antibody (Anti-Histone H3 (phospho S10) Abcam 47297 diluted 1:100 in block) at 4^°^C with rocking. The following morning the worms were washed 3X with PBS-Tween at 20^°^C with rocking for more than ten minutes. After washing, the worms were incubated with secondary antibody (Goat Anti-Rabbit IgG H&L (Alexa Fluor® 555) Abcam150078 diluted 1:1000 in PBS) for two hours at 20^°^C with rocking. The worms were washed again 3X with PBS-Tween as above and suspended in 15μL Vectashield with DAPI (Vector Laboratories, Burlingame, CA). A glass micropipet was used to transfer the worms in Vectashield to 2% agarose pads where they were covered with a glass coverslip for imaging.

### *C. elegans* staining for cell death

Animals were stained with SYTO12 (Invitrogen) following (Gumienny et al., 1999). This protocol requires the hermaphrodites to take up the dye and later, to consume unstained bacteria to clear the stain from the intestine. Since the worms are alive during the staining protocol, staining began in the morning and finished in the afternoon of Day 3 of adulthood.

### Staining and counting germline precursor cells

Hermaphrodites were aged in small populations of 30 per plate on either treatment or control plates. Unless indicated otherwise, Day 5 adult hermaphrodites were stained with DAPI (4′,6-diamidino-2-phenylindole) as described (Aprison and Ruvinsky, 2016) using a variation of the protocol by (Pepper et al., 2003).

Following washes in M9 and fixation with 95% ethanol, animals mounted in Vectashield, as above. We counted the number of nuclei in the proliferative zone, as defined by (Crittenden et al., 2006). In addition to the nuclei in mitotic cell cycle, this population contains some nuclei in the early stages of meiosis (Fox et al., 2011).

### Embryonic lethality measurements in mated *C. elegans* hermaphrodites

Synchronized populations of hermaphrodites were aged in small populations of ∼30 worms. On Day 5 of adulthood (when most of the worms were depleted of self-sperm), the hermaphrodites were singled and allowed to mate with 3 young adult N2 males for one hour. After mating, the hermaphrodites were singled to fresh plates. Progeny production was monitored for three days after the mating. The numbers of hatched progeny and unhatched fertilized eggs were scored.

### Male frequency experiments in *C. elegans* hermaphrodites

In selfing XX *C. elegans* hermaphrodites, rare male progeny (XO) arise due to chromosomal nondisjunction (Hodgkin et al., 1979), the incidence of which increases in older mothers (Luo et al., 2010; Rose and Baillie, 1979). To evaluate the frequency of male progeny, hermaphrodites were maintained in populations of 30 per plate (both control and treatment) and transferred every other day to fresh plates. On Day 4 of adulthood, the hermaphrodites were singled to fresh plates and the fraction of male offspring generated during the last two days of self-progeny production was noted.

### Germline morphology in aged *C. elegans* hermaphrodites

On Day 7 of adulthood, hermaphrodites were assessed for defects in the proximal gonad. To assess oocyte morphology, we used two phenotypes – abnormally small oocytes in the proximal gonad and the presence of atypical cavities between oocytes in the proximal gonad – as described (Luo et al., 2010). In young adult hermaphrodites, cellular volume of the oocytes in the proximal gonad increases prior to oocyte maturation so that the most proximal oocytes fill the gonadal lumen. Smaller oocytes have been associated with decreased oocyte quality (Andux and Ellis, 2008). Also prior to maturation, the oocyte shape changes from cylindrical to ovoid (McCarter et al., 1999). This shape change causes a small gap between the most proximal oocyte in the gonad and the next oocyte. Presence of cavities between more distal oocytes is abnormal. Presence of small oocytes or cavities in one arm of the gonad was scored as a “mild” defect, presence in both arms of the gonad was counted as “severe”.

### *D. melanogaster* egg chambers stages and immunohistochemistry

To produce synchronized populations of flies, virgin females were collected and crossed with young males. After 24 hours, males were removed and fertilized females were divided into two groups – one group was kept on control food, the other group on standard food supplemented with ∼0.9μM fluoxetine After two days flies were transferred to fresh food – control or conditioned with fluoxetine – and maintained an additional two days On Day 5 ovaries were dissected in ice-cold PBST (PBS with 0.5% Tween 20), individual ovarioles separated, fixed for 20 min with 4% paraformaldehyde, rinsed three times with PBS, incubated for 30 min with 0.3% BSA (blocking solution) in PBS at room temperature. Stages of egg chambers were determined as described (Jia et al., 2016). Follicle cells were counted in images that showed sagittal sections of each egg chamber. Incubation of samples with antibodies, rinses, and mounting for imaging were done as for *C. elegans* above.

### Embryonic lethality measurements in *D. melanogaster*

To produce synchronized populations of flies, virgin females were collected and crossed with young males. After 24 hours, males were removed and fertilized females were divided into two groups – one group was kept on control food, the other group on standard food supplemented with 0.9μM fluoxetine. Flies were maintained individually in 1.5 ml microcentrifuge tubes with a small hole made in the cap to allow air exchange with approximately 200 mg of food. Tubes were labelled and individual flies were followed throughout the entire experiment. Flies were transferred to fresh tubes every 24 hours. Tubes containing eggs were incubated 48 h and then examined to assess the number and state of the developing larvae. Every 10 days (Day 1, Day 10, and Day 20) two male flies were added to the tubes with female flies and allowed to mate for 24 hours. Flies and progeny production were followed for 30 days and embryonic lethality was determined for progeny produced in the final 15 days.

## Acknowledgements

We thank Rick Morimoto for generous hospitality and Bob Holmgren for flies. This work was funded in part by an NIH (R01GM126125) grant to IR. We thank WormBase and the Caenorhabditis Genetics Center (CGC). WormBase is supported by grant U41 HG002223 from the National Human Genome Research Institute at the NIH, the UK Medical Research Council, and the UK Biotechnology and Biological Sciences Research Council. The CGC is funded by the NIH Office of Research Infrastructure Programs (P40 OD010440).

