## Supplementary material for "Serotonergic signaling plays a deeply conserved role in improving oocyte quality": Figure S1

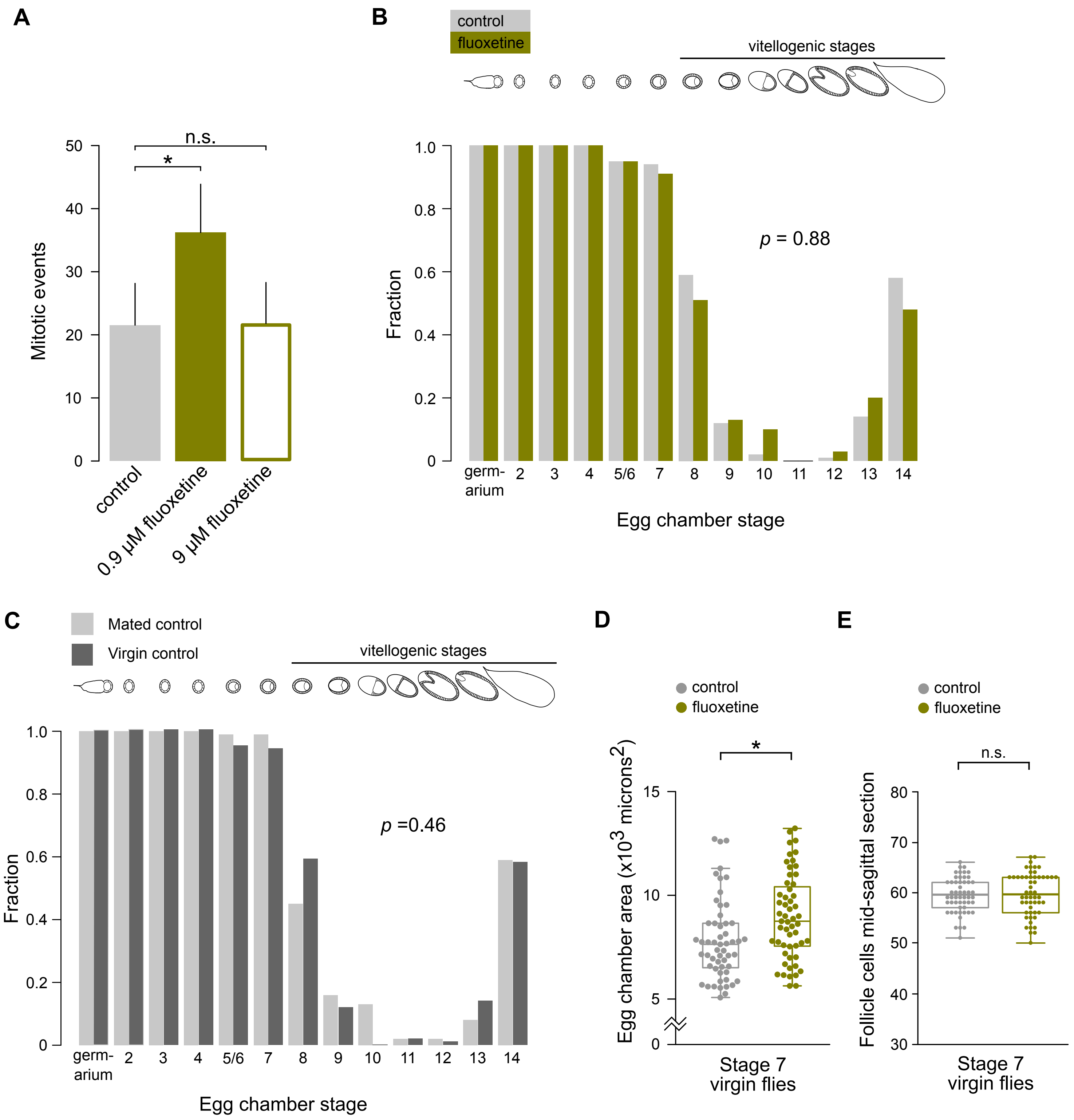
A
B
control
fluoxetine
vitellogenic stages
n.s.
*
Mitotic events
50
40
30
20
10
0
control
0.9 µM fluoxetine
9 µM fluoxetine
Fraction
1.0
0.8
0.6
0.4
0.2
0
p = 0.88
germ-arium
2
3
4
5/6
7
8
9
10
11
12
13
14
Egg chamber stage
C
Mated control
Virgin control
vitellogenic stages
p =0.46
D
E
control
fluoxetine
Egg chamber area (x10^3 microns^2)
15
10
5
0
Stage 7 virgin flies
Follicle cells mid-sagittal section
80
70
60
50
40
30

**Figure S1. Effects of fluoxetine at different concentration and in virgin females.**

(**A**) The number of mitotic events increased in flies raised on 0.9μM fluoxetine but not on 9μM. Error bars denote standard deviation. (**B**) Virgin flies (Day 5 of adulthood) on fluoxetine did not have more vitellogenic-stage egg chambers. **(C)** Mating alone did not alter the number of vitellogenic-stage egg chambers. Data from Figure 3A and Figure S1B. (**D**) Area of Stage 7 egg chambers in virgin Day 5 females raised on fluoxetine vs. untreated control. (**E**) The number of follicle cells in mid-sagittal sections of Stage 7 egg chambers in virgin Day 5 females raised fluoxetine vs. untreated control. *, $p<0.05$; **, $p<0.01$; ***, $p<0.001$. See Table S1 for primary data and details of statistical analyses.
